# Recurrent *FBXW7* mutations bypass Wnt/β-catenin addiction in cancer

**DOI:** 10.1101/2023.07.28.550933

**Authors:** Zheng Zhong, David M. Virshup

**Affiliations:** Program in Cancer and Stem Cell Biology, Duke-NUS Medical School, Singapore 169857; Department of Pediatrics, Duke University, Durham, North Carolina, USA 27710

## Abstract

Pathologic Wnt/β-catenin signaling drives various cancers, leading to multiple approaches to drug this pathway. Appropriate patient selection can maximize success of these interventions. Wnt ligand addiction is a druggable vulnerability in *RNF43*-mutant/*RSPO*-fusion cancers. However, pharmacologically targeting the biogenesis of Wnt ligands, e.g., with PORCN inhibitors, has shown mixed therapeutic responses, possibly due to tumor heterogeneity. Here we show that the tumor suppressor *FBXW7* is frequently mutated in *RNF43*-mutant/*RSPO*-fusion tumors, and *FBXW7* mutations cause intrinsic resistance to anti-Wnt therapies. Mechanistically, inactivation of FBXW7 stabilizes multiple oncoproteins including Cyclin E and MYC, and antagonizes the cytostatic effect of Wnt inhibitors. Moreover, although *FBXW7* mutations do not mitigate β-catenin degradation upon Wnt inhibition, *FBXW7*-mutant *RNF43*-mutant/*RSPO*-fusion cancers instead lose dependence on β-catenin signaling, accompanied by dedifferentiation and loss of lineage specificity. These *FBXW7*-mutant Wnt/β-catenin-independent tumors are susceptible to multi-CDK inhibition by dinaciclib. An in depth understanding of primary resistance to anti-Wnt/β-catenin therapies allows for more appropriate patient selection and use of alternative mechanism-based therapies.

## Introduction

Active Wnt signaling maintains the stem cell populations in multiple adult tissues while its hyperactivation due to genetic alterations drives many human cancers^1, 2^. Canonical Wnt signaling is activated by the engagement of secreted Wnt ligands with cell surface receptors Frizzleds (FZDs) and LRP5/6, leading to inactivation of GSK3 and stabilization of the transcriptional regulator β-catenin and other effector proteins such as MYC. Two categories of genetic events activate the Wnt pathway in human cancers: (1) downstream mutations, e.g., *APC* truncating mutations or *CTNNB1* (β-catenin) degron site mutations, that dissociate β-catenin regulation from upstream signaling and lead to its constitutive stabilization and activation; (2) mutations in the upstream regulators of Wnt receptors, such as inactivating mutations in the FZD-targeting E3 ubiquitin ligase genes *RNF43*/*ZNRF3* or gene fusions involving *RSPO2*/*3* leading to R-spondin overexpression that inactivates RNF43/ZNRF3^3, 4^. Both *RNF43*/*ZNRF3* mutations and *RSPO2*/*3* fusions up-regulate cell surface abundance of Wnt receptors and hypersensitize cells to Wnt ligands. These upstream mutations can be detected in diverse gastrointestinal cancers, e.g., *RNF43* mutations are present in 5-10% pancreatic cancers and *RSPO2*/*3* fusions are found in 2-10% colorectal cancers^4–9^.

Notably, while it remains challenging to therapeutically target stabilized β-catenin in cancers, the *RNF43*-mutant/*RSPO*-fusion cancers are attractive because preclinical studies revealed that multiple cancers with these molecular lesions are hypersensitive to upstream Wnt signaling blockade. Multiple modalities have been developed to achieve this and have had preclinical successes^3, 10, 11^. These include targeting the biogenesis of Wnt ligands with small molecule PORCN inhibitors^3, 10^, blocking the interaction between Wnt ligands and receptors using anti-FZD antibodies or recombinant Wnt decoy receptors^12–14^, and targeting R-spondins using RSPO2/3-neutralizing antibodies^11, 15^. Most notably, several PORCN inhibitors are being actively investigated in clinical trials, and it is anticipated that patient selection will be driven at least in part by these biomarkers^1^.

Notably, the therapeutic response to PORCN inhibitors is heterogeneous in *RNF43*-mutant/*RSPO*-fusion cancer patients^16, 17^. On one hand, durable responses were observed in several patients, confirming the important role of Wnt addiction and supporting the potential clinical benefits of PORCN inhibitors. On the other hand, intrinsic or acquired drug resistance was frequently observed during PORCN inhibitor treatment. However, the underlying mechanisms for PORCN inhibitor resistance remain elusive.

To identify potential drug resistance mechanisms, we previously performed an *in vivo* CRISPR gene knockout screen during PORCN inhibitor treatment in mice bearing *RNF43*-mutant pancreatic cancer xenografts. This screen identified gene candidates whose loss could cause Wnt independence and led to the discovery that defects in the p300/GATA6/differentiation axis can bypass the anti-differentiation role of Wnt signaling specifically in pancreatic cancer^18^. However, this does not appear to explain the drug resistance observed in those non-pancreatic cancer models, most notably in colorectal cancers. In the current study, we comprehensively investigate another hit from the screen, *FBXW7*, that encodes the substrate recognition component of a SCF E3 ubiquitin ligase complex. We find that *FBXW7* is frequently mutated in *RNF43*-mutant/*RSPO*-fusion cancers regardless of the tissue of origin. Inactivation of FBXW7 causes intrinsic resistance to Wnt inhibition by stabilizing oncoproteins including MYC and Cyclin E. Interestingly, *FBXW7* mutants lose dependence on β-catenin, accompanied by dedifferentiation and loss of lineage specificity, suggesting a cancer evolutionary trajectory from Wnt-dependent low-grade tumors to Wnt-independent and dedifferentiated cancers. *FBXW7* mutations may be a powerful biomarker for patient selection, as they may drive a major drug resistance mechanism limiting the effectiveness of diverse therapies targeting Wnt/β-catenin signaling.

## Results

### Recurrent *FBXW7* mutations in *RNF43*-mutant/*RSPO*-fusion cancers

Our previous *in vivo* CRISPR screen identified several genes whose knockout caused resistance to a PORCN inhibitor^18^. Most of these genes are known negative regulators of Wnt pathway, e.g., *APC* and *AXIN1*, so that their knockout provides downstream activation of Wnt/β-catenin signaling. These genes were independently observed in a genome-wide *in vitro* screen performed by the Angers group aiming to find resistance genes against another PORCN inhibitor^19^. One unexpected hit from both screens was *FBXW7*, a known tumor suppressor but one that is not obviously involved in regulating Wnt signaling^20^. To study how alterations in these putative resistance genes might affect the overall clinical response of Wnt-targeting therapies, we analyzed their mutation frequency in *RNF43*-mutant/*RSPO*-fusion cancers. We focused on all types of genetic alterations of *FBXW7* and established Wnt/β-catenin-activating mutations including *APC* or *AXIN1* truncation/deletion and *CTNNB1* stabilizing missense mutations.

In a combined cohort of colorectal cancers harboring *PTPRK*-*RSPO3* fusions, the most common *RSPO*-fusion cancer type, *FBXW7* mutations were detected in 25% cases (**Fig. 1A**). Half of these *FBXW7* mutations were hotspot missense mutations that are known to be dominant-negative. Truncations of *FBXW7*, that are presumably inactivating, were also common. This mutational landscape is similar to the *FBXW7* mutation landscape in the bulk of colorectal cancers, largely due to *APC* mutations^20^. In contrast, mutations in *APC*, *AXIN1*, and *CTNNB1* were very rare in *RSPO*-fusion colorectal cancers, as only one *APC* truncation was detected (**Fig. 1A**).

**Figure 1.**
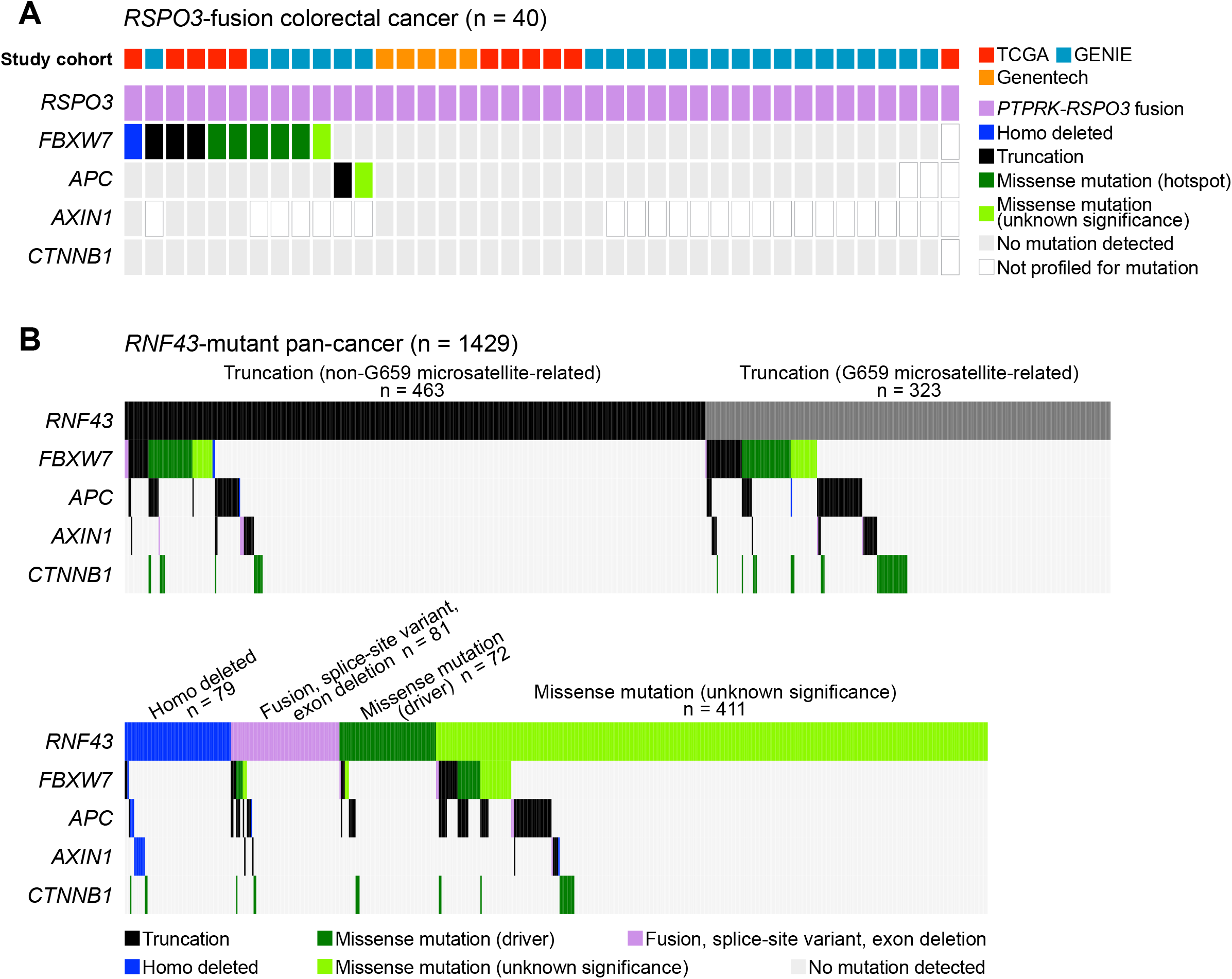
Recurrent *FBXW7* mutations in *RNF43*-mutant/*RSPO*-fusion cancers. **(A)** Recurrent *FBXW7* mutations in 10 (25%) individuals in a combined cohort of 40 *PTPRK*-*RSPO3* fusion colorectal tumors. Each column represents an individual tumor sample. **(B)** Mutations in *FBXW7*, *APC*, *AXIN1*, and *CTNNB1* in 1429 *RNF43*-mutant tumors regardless of tumor origin (pan-cancer). Each column represents an individual tumor sample, grouped based on the type of *RNF43* mutations. For *APC*, *AXIN1*, and *CTNNB1*, only established Wnt/β-catenin-activating mutations such as *APC*/*AXIN1* truncation/deletion and hotspot *CTNNB1* stabilizing mutations are shown, while missense mutations that are empirically recognized as probable wildtype-like passenger mutations are not shown.

*RNF43* mutations appear in multiple cancer types, so we performed a pan-cancer analysis of *RNF43* and those putative drug resistance genes (**Fig. 1B**). As the role of the recurrent *RNF43* G659 microsatellite-related frameshift mutations remains controversial^21–24^, we analyzed those separately from the non-G659-related truncations. For missense mutations of *RNF43*, we annotated them based on previous characterization studies^23, 25^ and classified them as driver mutations (loss of function, partial loss of function, or dominant-negative) or mutations of unknown significance. Cancer cases with known benign missense mutations in *RNF43* were excluded. Importantly, *FBXW7* mutations were detected in 16.7% of all *RNF43*-mutant cancers. Again, around half of these *FBXW7* alterations were hotspot missense mutations. Notably, different from *RSPO*-fusion colorectal cancers, a significant subset (17.8%) of *RNF43*-mutant cancers harbored downstream β-catenin-stabilizing mutations in either *APC*, *AXIN1*, or *CTNNB1*.

In summary, we identified recurrent *FBXW7* mutations in 16-25% of both *RNF43*-mutant and *RSPO3*-fusion cancers. In *RNF43*-mutant cancers, we also observed frequent *APC*, *AXIN1*, or *CTNNB1* mutations that stabilize and activate β-catenin. These mutations could result in resistance to anti-Wnt therapies in *RNF43*-mutant/*RSPO3*-fusion cancers.

### *FBXW7* mutation status predicts resistance to Wnt inhibition

To evaluate if pre-existing *FBXW7* mutations predicted resistance to anti-Wnt therapies, we collected a panel of cancer cell lines from diverse lineages harboring inactivating mutations in *RNF43*/*ZNRF3* or *RSPO2*/*3* fusions and tested their response to the PORCN inhibitor ETC-159 in a soft agar colony formation assay (**Fig. 2A**). As previously reported^3, 26, 27^, HPAF-II and other cell lines without other Wnt pathway mutations failed to form colonies in the presence of the PORCN inhibitor. As expected, HuG1-N and HCT116, the two cell lines with additional APC truncation and β-catenin-stabilizing mutation, respectively, were resistant to ETC-159. Importantly, the three cell lines harboring *FBXW7* mutations, AsPC-1 and NCIH2172 with hotspot mutations and HuCCT1 with a truncating mutation, were either less sensitive or highly resistant to ETC-159.

**Figure 2.**
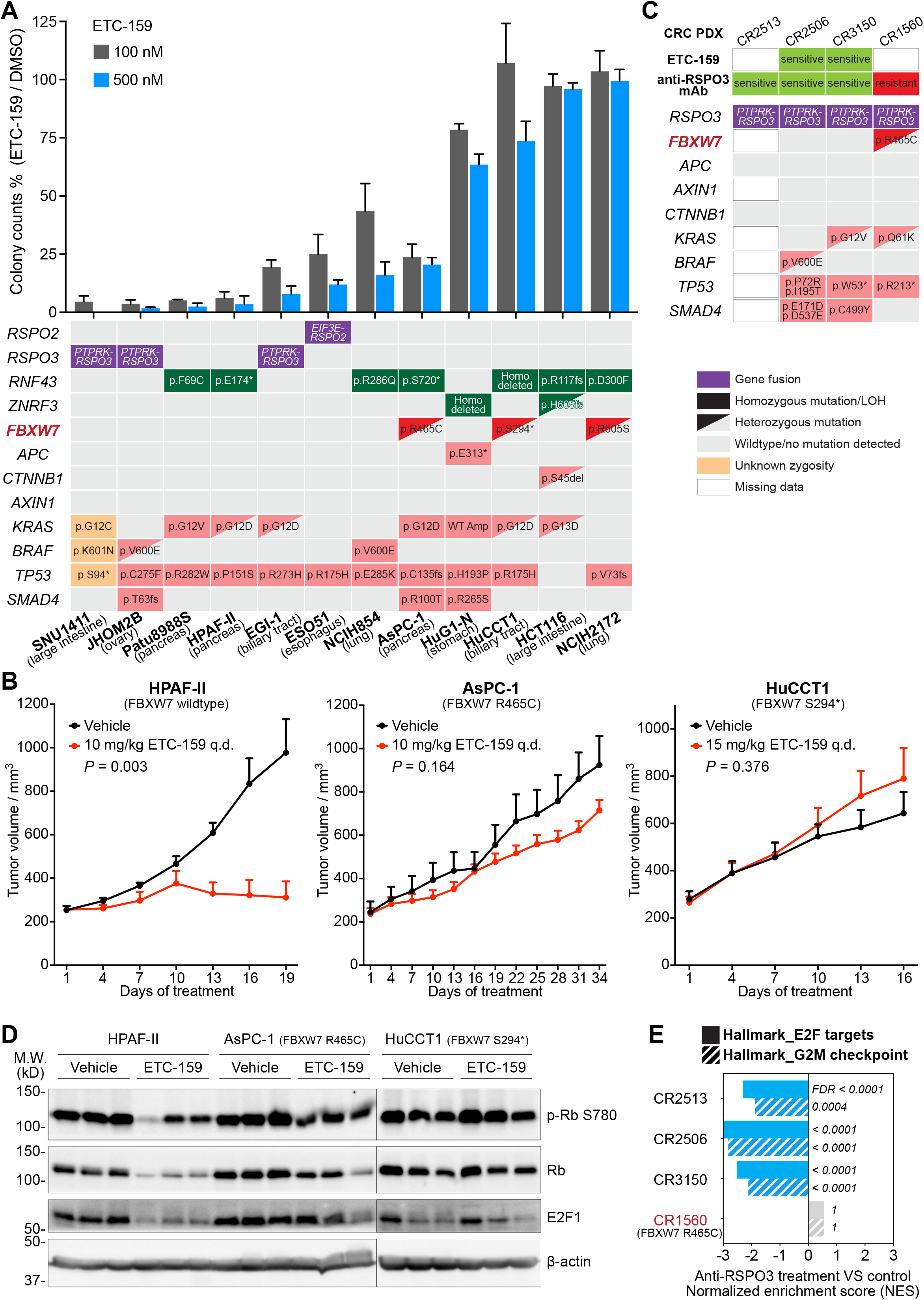
*FBXW7* mutation status predicts resistance to Wnt inhibition in *RNF43*-mutant/*RSPO*-fusion cancers. **(A)** *RNF43*-mutant/*RSPO*-fusion cancer cell lines harboring inactivating mutations in *FBXW7* were resistant to PORCN inhibitor in *in vitro* colony formation assays. Cancer cells seeded in soft agar were treated with 100 or 500 nM PORCN inhibitor ETC-159 or DMSO (0.1% v/v) for 1-2 weeks. JHOM2B and HuCCT1 cells did not grow in soft agar and were examined in a low-density 2D culture assay. Colonies were quantified and data are plotted as mean + SD reflecting the relative colony formation (ETC-159-treated versus DMSO control). n = 2-4 biological replicates per condition. Genetic alterations in drug sensitizing genes (*RNF43*, *ZNRF3*, and *RSPO2*/*3*), drug resistance genes (*FBXW7*, *APC*, *AXIN1*, and *CTNNB1*), and additional oncogenic drivers are shown in the lower panel. **(B)** *RNF43*/*FBXW7*-double mutant tumors were resistant to PORCN inhibitor *in vivo*. HPAF-II and AsPC-1 pancreatic cancer cells and HuCCT1 cholangiocarcinoma cells were subcutaneously injected into NSG mice. After tumor establishment, tumor-bearing mice were treated with vehicle or 10 or 15 mg/kg ETC-159 once every day (q.d.). Data are represented as mean + SEM (n = 6-8 tumors/arm). *P* values of 2-tailed, unpaired *t* test between vehicle and ETC-159 arms at the last time point are shown. **(C)** An *RSPO3*-fusion colorectal cancer PDX resistant to RSPO3-neutralizing antibody harbors *FBXW7* hotspot mutation. Sensitivity of these *RSPO3*-fusion colorectal cancer PDXs to ETC-159 and the anti-RSPO3 monoclonal antibody (mAb) was reported in ref. 15. **(D, E)** Wnt inhibition failed to induce cytostasis in *RNF43*-mutant/*RSPO*-fusion tumors harboring *FBXW7* mutations. **(D)** Tumors harvested from **(B)** were analysed by western blotting for cell cycle-related proteins. Each lane represents an independent tumor. **(E)** Gene set enrichment analysis (GSEA) of cell cycle-related hallmark gene sets for *RSPO3*-fusion PDXs mentioned in **(C)** treated with the RSPO3-neutralizing mAb versus control (ref. 15). The transcriptomic data were extracted from GSE73906.

To test if these *FBXW7* mutations affected drug sensitivity *in vivo*, we examined the growth inhibition efficacy of ETC-159 in mouse xenografts of three pancreatobiliary cancer cell lines with *RNF43* mutations (**Fig. 2B**). Consistent with the *in vitro* observation, while ETC-159 potently suppressed the *FBXW7*-wildtype HPAF-II tumor growth, at the same dose the AsPC-1 tumor had only 30% tumor growth inhibition, and there was no effect on the HuCCT1 tumor.

In a preclinical study from OncoMed using an RSPO3-neutralizing antibody to treat *RSPO3*-fusion colorectal cancers^15^, three of four *RSPO3*-fusion positive patient-derived xenografts (PDXs) showed tumor growth inhibition, while a fourth tumor, CR1560, was unaffected. We speculated that this resistance could be due to a pre-existing *FBXW7* mutation. Indeed, inspection of mutation profiles of these four PDXs revealed a hotspot p.R465C mutation of *FBXW7* only in the drug resistant PDX (**Fig. 2C**). Moreover, in these PDXs and the above tested cancer cell lines, the sensitivity to anti-Wnt therapies was not related to the other oncogenic driver mutations including alterations in *KRAS*, *BRAF*, *TP53*, and *SMAD4* (**Fig. 2A** and **C**). These results, taken together, suggest that pre-existing *FBXW7* mutations are common in *RNF43*-mutant/*RSPO3*-fusion cancers and cause resistance to Wnt inhibitors.

Wnt inhibition leads to cell cycle arrest and differentiation rather than cell death in Wnt-addicted cancers^14, 18, 26, 28^. We examined if the molecular response to anti-Wnt therapies differed in the *FBXW7*-wildtype versus mutant tumors (**Fig. 2D**). In the *FBXW7*-wildtype HPAF-II xenografts, PORCN inhibition led to significant dephosphorylation of Rb protein and (as dephosphorylated Rb is less stable^29, 30^) reduced total Rb abundance. PORCN inhibition also markedly reduced the abundance of E2F1, a key regulator of the cell cycle progression. These indicate a potent cell cycle arrest in HPAF-II tumors, consistent with our prior studies^28^. In contrast, these effects were largely attenuated or even absent in the *FBXW7*-mutant AsPC-1 and HuCCT1 tumors (**Fig. 2D**). Similar results were noted when we reanalyzed the data from the OncoMed study of *RSPO3*-fusion colorectal cancer PDXs^15^, where the RSPO3-neutralizing antibody treatment led to potent cell cycle arrest in the three sensitive models but showed no effect in the *FBXW7*-mutant drug resistant PDX (**Fig. 2E**). Therefore, the association of *FBXW7* mutations and PORCN inhibitor resistance was confirmed on the molecular level in both *RNF43*-mutant and *RSPO*-fusion cancers.

Taken together, these results establish a strong correlation between *FBXW7* mutations and resistance to anti-Wnt therapies in *RNF43*-mutant/*RSPO*-fusion cancers. These also suggest that *FBXW7* mutation status can be used as a biomarker for patient selection for anti-Wnt therapies.

### Inactivation of FBXW7 confers intrinsic resistance to Wnt inhibition

To directly test if *FBXW7* mutations cause the drug resistance and to study the underlying mechanism, we inactivated *FBXW7* in PORCN inhibitor-sensitive cancer cells and examined if they became inhibitor resistant. Specifically, we truncated the endogenous FBXW7 protein approximately at amino acid position 420 in the *FBXW7-*wildtype *RNF43*-mutant HPAF-II pancreatic cancer cells using CRISPR/Cas9 (**Fig. S1**) and subsequently tested the sensitivity of the xenografts to PORCN inhibitor ETC-159 (**Fig. 3A**). Notably, immediately after initiation of this animal study, Singapore entered the unexpected COVID-19 lockdown period, during which time we only had limited access to the vivarium. Therefore, these tumor-bearing mice were treated with an intermittent drug dosing schedule that nevertheless produced a significant tumor growth inhibition in control tumors (**Fig. 3A**, left). In contrast, ETC-159 had no effect on the growth of tumors with FBXW7 truncations (**Fig. 3A**, right).

**Figure 3.**
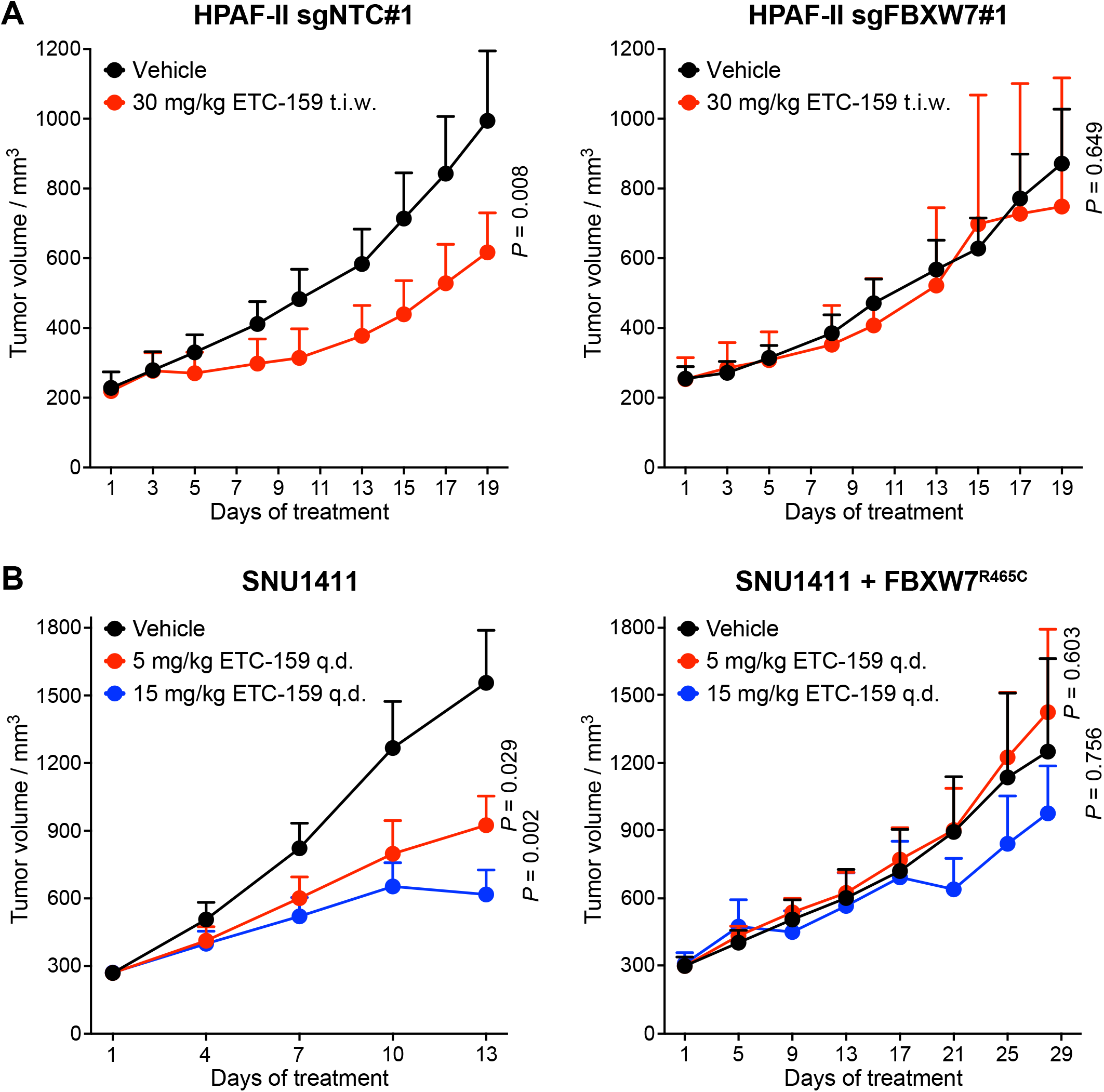
Inactivation of FBXW7 confers intrinsic resistance to Wnt inhibition in *RNF43*-mutant/*RSPO*-fusion cancers. **(A)** Truncating the endogenous FBXW7 in *RNF43*-mutant HPAF-II tumors caused resistance to ETC-159. HPAF-II cells transduced with a non-targeting control sgRNA (sgNTC#1) or a sgRNA targeting *FBXW7* (sgFBXW7#1) were subcutaneously injected into NSG mice. After tumor establishment, tumor-bearing mice were treated with vehicle or 30 mg/kg ETC-159 three times a week (t.i.w). Data are represented as mean + SEM (n = 6-7 tumors/arm). **(B)** Ectopic expression of FBXW7 p.R465C mutant (a hotspot dominant-negative mutation) in *RSPO3*-fusion SNU1411 tumors caused resistance to PORCN inhibition. SNU1411 cells or cells stably expressing the FBXW7 mutant were subcutaneously injected into NSG mice. After tumor establishment, mice were treated with vehicle or 5 or 15 mg/kg ETC-159 once every day (q.d.). Data are represented as mean + SEM (n = 6-10 tumors/arm). *P* values of 2-tailed, unpaired *t* test between vehicle and ETC-159 arms at the last time point are shown.

Half of the *FBXW7* mutations detected in clinical samples are hotspot missense mutations that known to be dominant-negative^20^ (**Fig. 1**). To examine the effect of these dominant-negative mutations on drug resistance, we stably expressed an exogenous FBXW7 p.R465C hotspot mutant in *FBXW7*-wildtype *RSPO3*-fusion SNU1411 colorectal cancer cells (**Fig. 3B**). Similar to the truncation mutants, while ETC-159 inhibited growth of the parental SNU1411 xenografts in a dose-dependent manner, the growth of tumors expressing the FBXW7 dominant negative mutant no longer responded to ETC-159 treatment (**Fig. 3B**). Taken together, these results demonstrate that inactivation of FBXW7 through either truncations or dominant-negative missense mutations confers intrinsic resistance to Wnt inhibition in *RNF43*-mutant/*RSPO*-fusion cancers.

### FBXW7 mutations function in part by stabilizing MYC and Cyclin E

We next investigated the molecular mechanisms underlying the drug resistance caused by FBXW7 inactivation. Previously we showed that in drug-sensitive *RNF43*-mutant/*RSPO*-fusion tumors Wnt inhibition down-regulated the cell cycle machinery due to suppression of the Wnt/β-catenin/MYC network^28^, and this led to a potent cell cycle arrest. As the substrate recognition subunit of a SCF E3 ubiquitin ligase complex, FBXW7 controls the degradation of multiple oncoproteins^20, 31^. We hypothesized that inactivation of FBXW7 leads to the accumulation or prevents the degradation of certain proteins that antagonize the cytostatic effect of Wnt inhibitors. We therefore examined the protein abundance of various known FBXW7 substrates in the HPAF-II and SNU1411 xenograft tumors treated with or without ETC-159. Out of the six reported FBXW7 substrates (**Fig. 4A**), we observed consistent up-regulation of Cyclin E1 in both HPAF-II and SNU1411 tumors after FBXW7 inactivation. In HPAF-II tumors, FBXW7 inactivation increased basal MYC abundance, while this effect was less evident in SNU1411 tumors. However, in both models, FBXW7 inactivation mitigated the degradation of MYC upon ETC-159 treatment (**Fig. 4A**). This was due to increased MYC stability after FBXW7 inactivation (**Fig. S2**). Other known FBXW7 substrates were regulated in a context-dependent manner, as we observed an increase in KLF5 abundance after FBXW7 inactivation in HPAF-II tumors, and an increase in PKM2 abundance only in SNU1411 tumors (**Fig. 4A**). Two other presumed FBXW7 substrates, c-Jun and the NOTCH1 intracellular domain, were not affected in either model.

**Figure 4.**
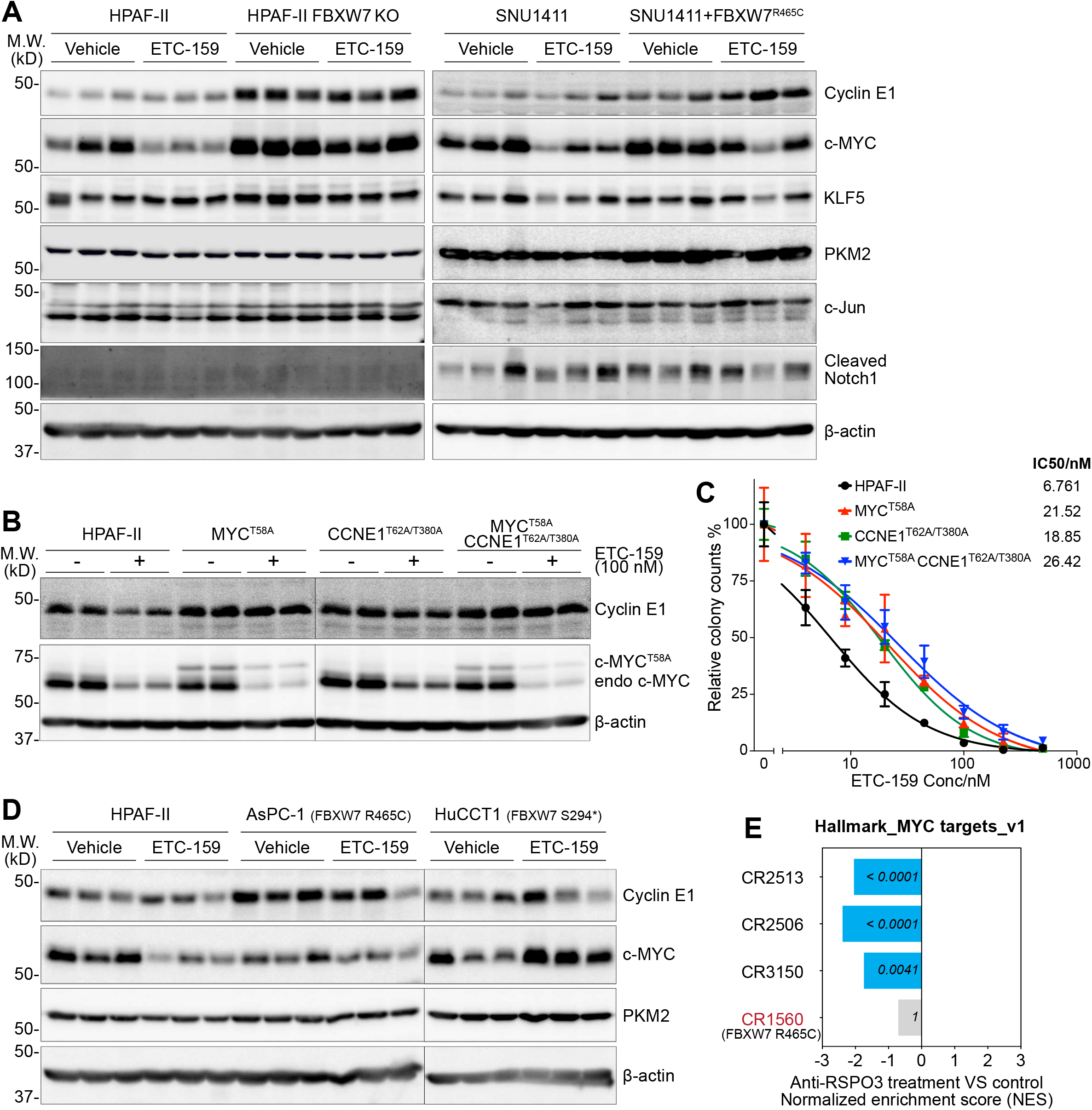
Stabilization of MYC and Cyclin E after FBXW7 inactivation causes resistance to Wnt inhibition. **(A)** *RNF43*-mutant/*RSPO*-fusion cancer xenografts **(Fig. 3)** were analysed by western blotting for 6 reported substrates of FBXW7. β-actin was used as loading control. Each lane represents an independent tumor. In the panel of SNU1411 ± FBXW7 mutant, ETC-159-treated samples were from the 5 mg/kg group shown in Fig. 3B. **(B, C)** Ectopic expression of human phosphorylation site mutant Cyclin E1 (CCNE1) or MYC that are resistant to GSK3/FBXW7-mediated degradation caused resistance to PORCN inhibition in HPAF-II cells. **(B)** HPAF-II cells or cells stably expressing the mutants were treated with DMSO or 100 nM ETC-159 for 4 days, followed by western blotting analysis. The exogenous MYC mutant was fused with 3xHA tag that led to the mobility shift on the gel. endo, endogenous. n = 2 biological replicates/condition. **(C)** HPAF-II cells or cells stably expressing the mutants were seeded in soft agar and treated with the indicated concentrations of ETC-159 for ∼2 weeks. Colony numbers were quantified and normalized to the DMSO control. Data are represented as mean ± SD (n = 3 biological replicates/condition). IC50 values are shown. **(D)** *RNF43*-mutant tumors with or without pre-existing *FBXW7* mutations and treated with vehicle or ETC-159 **(Fig. 2B)** were analysed by western blotting. Each lane represents an independent tumor. **(E)** GSEA of the hallmark “MYC targets” gene set for *RSPO3*-fusion PDXs **(Fig. 2C)** treated with the RSPO3-neutralizing mAb versus control. The transcriptomic data were extracted from GSE73906.

Stabilization of Cyclin E1 and MYC by FBXW7 inactivation was consistently observed in two independent models. To test whether these proteins mediated the FBXW7 inactivation-caused drug resistance, we ectopically expressed human mutant Cyclin E1 and/or MYC in HPAF-II cells (**Fig. 4B**). These mutants cannot be phosphorylated by GSK3 and so they cannot be subsequently recognized by FBXW7. Notably, expressing either mutant Cyclin E or mutant MYC led to a clear shift of the ETC-159 dose-response curve in the soft agar colony formation assay (**Fig. 4C**), reflecting drug resistance. Interestingly, co-expressing stabilized Cyclin E and MYC only modestly further enhanced the drug resistance (**Fig. 4C**), suggesting that Cyclin E and MYC might share redundant roles in this setting. This is consistent with the observation that expressing mutant MYC alone also up-regulated Cyclin E abundance (**Fig. 4B**). Thus, FBXW7 inactivation causes intrinsic resistance to Wnt inhibition at least partially due to stabilization of Cyclin E and MYC.

We extended this observation to the naturally pre-existing *FBXW7*-mutant drug resistant models. For example, the *FBXW7*-mutant AsPC-1 tumors had a significantly higher Cyclin E1 protein (but not mRNA) abundance than the *FBXW7*-wildtype HPAF-II tumors (**Fig. 4D** and **Fig. S3**). Moreover, in both *FBXW7*-mutant AsPC-1 and HuCCT1 tumors, ETC-159 treatment no longer led to a significant decrease of MYC protein (**Fig. 4D**). This is consistent with our reanalysis of the transcriptomic response data reported in the *RSPO3*-fusion colorectal cancer PDXs^15^, where treatment with RSPO3-neutralizing antibody significantly down-regulated the mRNA levels of MYC target genes in all the models except CR1560 bearing a hotspot *FBXW7* mutation (**Fig. 4E**). The above findings demonstrate that inactivation of FBXW7 confers intrinsic resistance to Wnt inhibition in *RNF43*-mutant/*RSPO*-fusion cancers at least in part through stabilizing MYC and Cyclin E.

### *FBXW7*-mutant *RNF43*-mutant/*RSPO*-fusion cancers lose their dependence on β-catenin signaling

*RNF43*-mutant/*RSPO*-fusion cancers are addicted to β-catenin signaling^3^. The resistance conferred by *FBXW7* mutations functions through stabilization of MYC and Cyclin E. Because β-catenin is regulated by a different E3 ubiquitin ligase that uses β-TrCP as a substrate recognition subunit^32^, *FBXW7* mutations should have no effect on β-catenin degradation upon Wnt inhibition (**Fig. 5A**). This raised an interesting question – was β-catenin stable in *FBXW7*-mutant cancers, or did the *FBXW7*-mutant cancers no longer rely on β-catenin?

**Figure 5.**
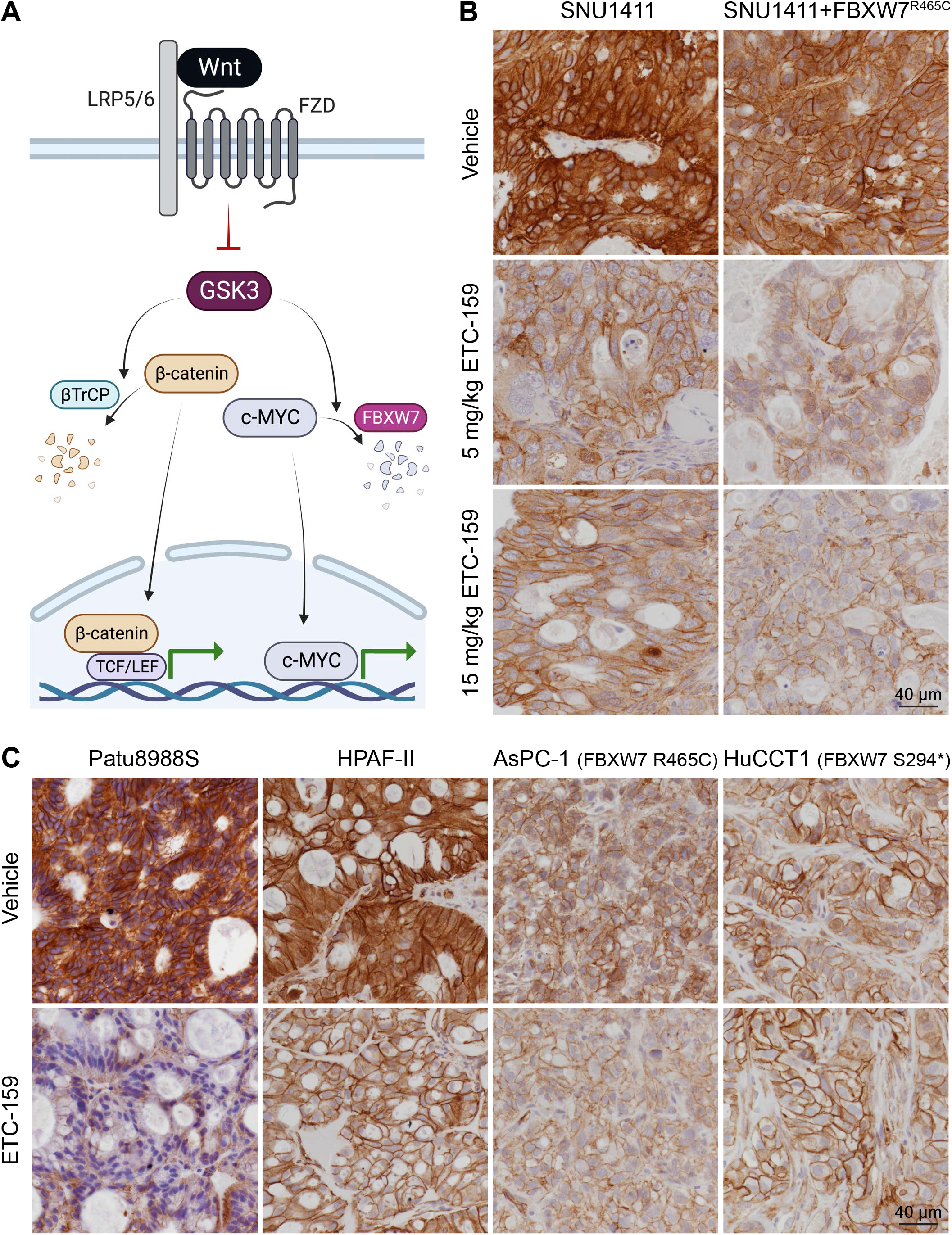
Inactivation of FBXW7 does not mitigate β-catenin degradation upon Wnt inhibition and *FBXW7*-mutant *RNF43*-mutant/*RSPO*-fusion cancers show low basal levels of β-catenin. **(A)** Schematic model of the Wnt/GSK3-regulated β-catenin and MYC turnover. Please refer to the text for detailed description. **(B)** PORCN inhibition decreased β-catenin abundance in *RSPO3*-fusion SNU1411 tumors regardless of FBXW7 status. β-catenin staining was performed on SNU1411 tumor samples with or without expressing the dominant-negative FBXW7 mutant and treated with vehicle or ETC-159 **(Fig. 3B)**. **(C)** Pre-existing *RNF43*/*FBXW7*-double mutant cancer xenografts showed low basal levels of β-catenin in both cytoplasm and nucleus. β-catenin staining was performed on tumor samples harvested from experiments shown in Fig. 2B and our previous study (for the *RNF43*-mutant *FBXW7*-wildtype Patu8988S pancreatic cancer xenografts, ref. 18). Note that the β-catenin staining on HPAF-II, AsPC-1, and HuCCT1 samples was performed in the same batch while Patu8988S samples were assayed in another batch.

To test if FBXW7 inactivation affects β-catenin degradation upon Wnt inhibition, we used immunohistochemistry to examine β-catenin abundance in the *RSPO3*-fusion SNU1411 colorectal tumor models. As expected, the basal β-catenin level was high in both cytoplasm and nucleus of the xenograft tumors, and PORCN inhibition robustly reduced β-catenin abundance (**Fig. 5B**). Importantly, the reduction of β-catenin was similarly observed in xenografts that expressed the dominant-negative FBXW7 mutant (**Fig. 5B**), confirming that inactivation of FBXW7 does not mitigate β-catenin degradation. Since these *FBXW7*-mutant SNU1411 tumors continued growing during ETC-159 treatment (**Fig. 3B**), they must have lost their initial dependence on β-catenin. We extended this observation in additional *RNF43*-mutant tumor xenografts. As expected, *FBXW7*-wildtype tumors had high basal levels of β-catenin that significantly decreased after Wnt inhibition (**Fig 5C**, left). Notably, the *FBXW7*-mutant tumors showed lower levels of β-catenin in both the cytoplasm and nucleus even without ETC-159 treatment (**Fig. 5C**, right). These observations suggest that during cancer evolution *RNF43*-mutant/*RSPO*-fusion cancer cells lose their dependence on Wnt/β-catenin after acquiring *FBXW7* mutations.

To test if β-catenin indeed becomes dispensable, we explored the specific requirements for β-catenin and MYC in *FBXW7* wildtype and mutant xenograft tumors. We used a dCas9-KRAB-mediated gene knockdown system where the expression of sgRNAs targeting the *CTNNB1* or *MYC* promoters could be induced by doxycycline treatment. As expected, knockdown of β-catenin in the Wnt-dependent HPAF-II cells potently suppressed tumor growth (**Fig. 6A** and **B**). As *MYC* is a β-catenin target gene, β-catenin knockdown in HPAF-II cells partially reduced *MYC* mRNA abundance (**Fig. 6A**). In contrast, direct knockdown of *MYC* transcription had a much smaller effect on HPAF-II tumor growth and did not phenocopy the tumor growth inhibition caused by β-catenin knockdown (**Fig. 6A** and **B**). This is consistent with our understanding that in Wnt-addicted cancers β-catenin plays more roles besides transcriptionally regulating *MYC* expression^28^.

**Figure 6.**
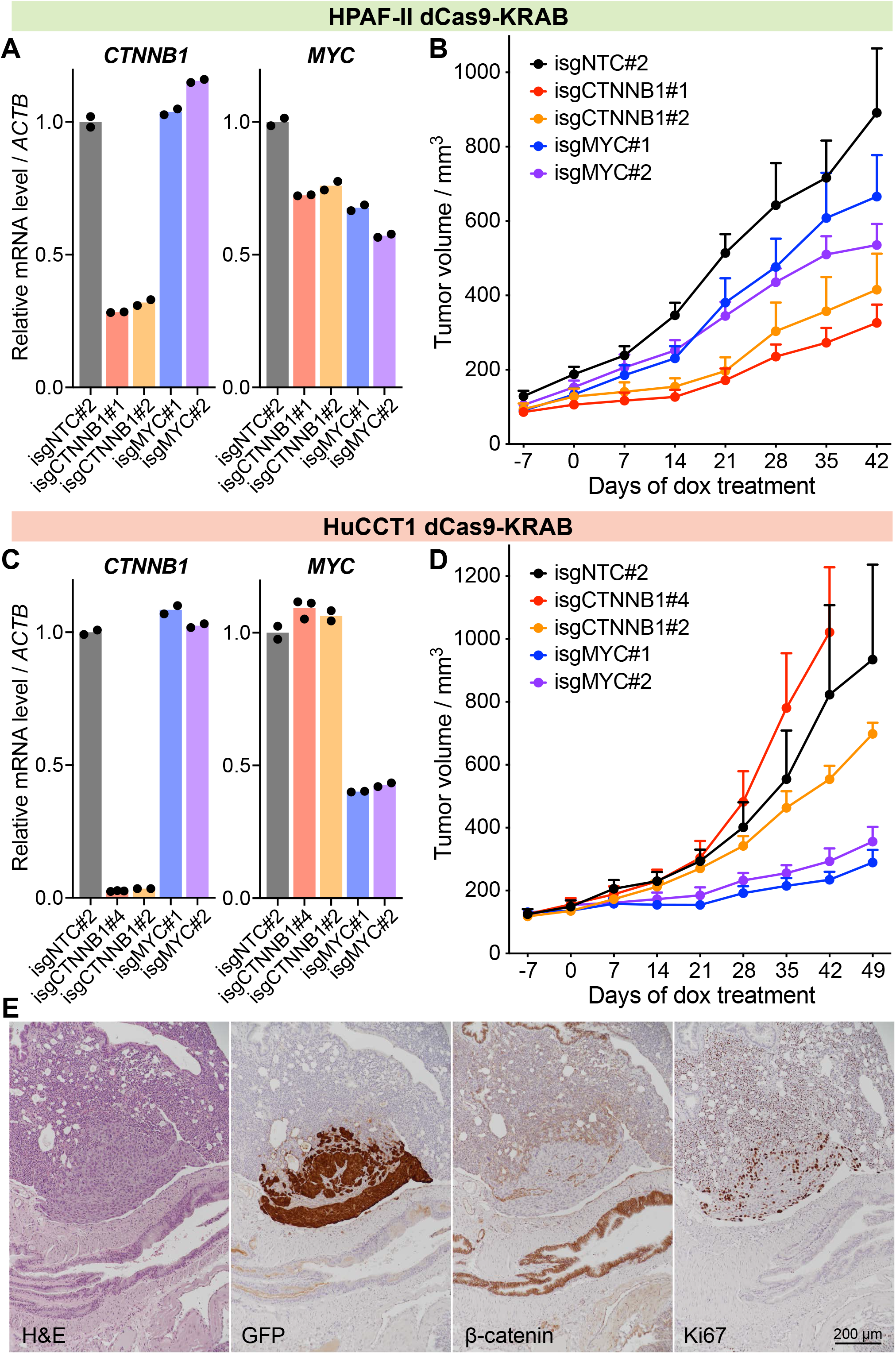
*RNF43*/*FBXW7*-double mutant cancer cells lost dependence on β-catenin. **(A, B)** Knockdown of *CTNNB1* or *MYC* in *FBXW7*-wildtype HPAF-II cells and the effects on tumor growth. **(A)** HPAF-II cells stably expressing dCas9-KRAB were transduced with a doxycycline-inducible sgRNA expression vector targeting *CTNNB1* or *MYC* promoters with two independent sgRNAs per gene, or a non-targeting control (isgNTC#2). These cells were treated with doxycycline for 4 days, followed by RT-qPCR analysis. **(B)** These cells without doxycycline treatment were subcutaneously injected into NSG mice. After tumor establishment, mice were put on continuous doxycycline feed until the end of the study. Data are represented as mean + SEM (n = 6 tumors/arm). **(C, D)** Knockdown of *CTNNB1* or *MYC* in *FBXW7*-mutant HuCCT1 cells and the effects on tumor growth. Experiments were performed as in **(A, B)**. **(D)** Data are represented as mean + SEM (n = 6-8 tumors/arm). **(E)** HuCCT1 cells retained spontaneous metastatic capability after β-catenin knockdown. Representative images of H&E, GFP, β-catenin, and Ki67 staining of a lung metastasis (out of more than 10 metastatic lesions per lung section) from a mouse with the HuCCT1 isgCTNNB1#4 subcutaneous tumor. The sgRNA-expressing vector also expressed GFP, which was stained to distinguish the cancer cells from the host tissues.

In sharp contrast, in the *RNF43*-deleted but *FBXW7*-mutant HuCCT1 xenografts, depleting β-catenin no longer affected *MYC* transcription nor tumor growth (**Fig. 6C** and **D**). Instead, knockdown of *MYC* potently inhibited HuCCT1 tumor growth (**Fig. 6D**), suggesting that these *FBXW7*-mutant cancer cells had lost dependence on β-catenin and had shifted their oncogene addiction to MYC. Moreover, β-catenin knockdown did not affect the spontaneous lung metastasis occurred in HuCCT1 xenograft-bearing mice. Histopathologic analysis confirmed that cancer cells of the lung metastases were highly proliferative despite the absence of β-catenin (**Fig. 6E**), confirming that β-catenin is dispensable in this model. Similarly, we knocked down β-catenin in AsPC-1 xenografts that harbor a *FBXW7* mutation but partially responded to Wnt inhibition. We speculate that these tumors may represent a transition state towards Wnt/β-catenin independence. β-catenin knockdown delayed initial AsPC-1 tumor growth, but after a period of adaptation these tumors grew as fast as control (**Fig. S4A**). As with the HuCCT1 tumors, *MYC* transcription was no longer affected by β-catenin knockdown in these AsPC-1 xenograft tumors (**Fig. S4B**), consistent with the loss of β-catenin dependence. In summary, the data indicate that FBXW7 is not directly involved in the Wnt/β-catenin axis, but that *RNF43*-mutant/*RSPO*-fusion cancers lose their dependence on β-catenin signaling after acquiring *FBXW7* mutations.

### Inactivation of FBXW7 leads to dedifferentiation and phenotypic switching in *RNF43*-mutant/*RSPO*-fusion cancers

Hyperactivated Wnt/β-catenin signaling in *RNF43*-mutant/*RSPO*-fusion tumors maintains cancer stemness, while Wnt inhibition promotes cancer cell differentiation^10, 11, 18^. Since *FBXW7*-mutant *RNF43*-mutant/*RSPO*-fusion cancers have low levels of β-catenin (**Fig. 5C**), we speculated that these cancers might be more differentiated. However, while cancer cells in the *FBXW7*-wildtype HPAF-II tumors formed large glandular-like epithelial structures, reflecting a well/moderately differentiated state, these structures were rarely seen in the *FBXW7*-mutant tumors (**Fig. 7A**). Consistent with a more mesenchymal phenotype, these *FBXW7*-mutant cancer cells have low expression of the epithelial marker E-cadherin and high expression of the mesenchymal marker vimentin (**Fig. S5**). We further examined the expression of mucin as a marker of differentiation in these xenografts. The *FBXW7*-mutant tumors showed comparable if not less mucin staining compared with the *FBXW7*-wildtype HPAF-II tumors (**Fig. 7A**). Notably, while Wnt inhibition significantly promoted mucin expression in HPAF-II tumors, it had no effect in the two *FBXW7*-mutant cancers (**Fig. 7A**). In conclusion, *FBXW7*-mutant *RNF43*-mutant/*RSPO*-fusion tumors appeared to be less differentiated and are refractory to the Wnt inhibition-induced differentiation program.

**Figure 7.**
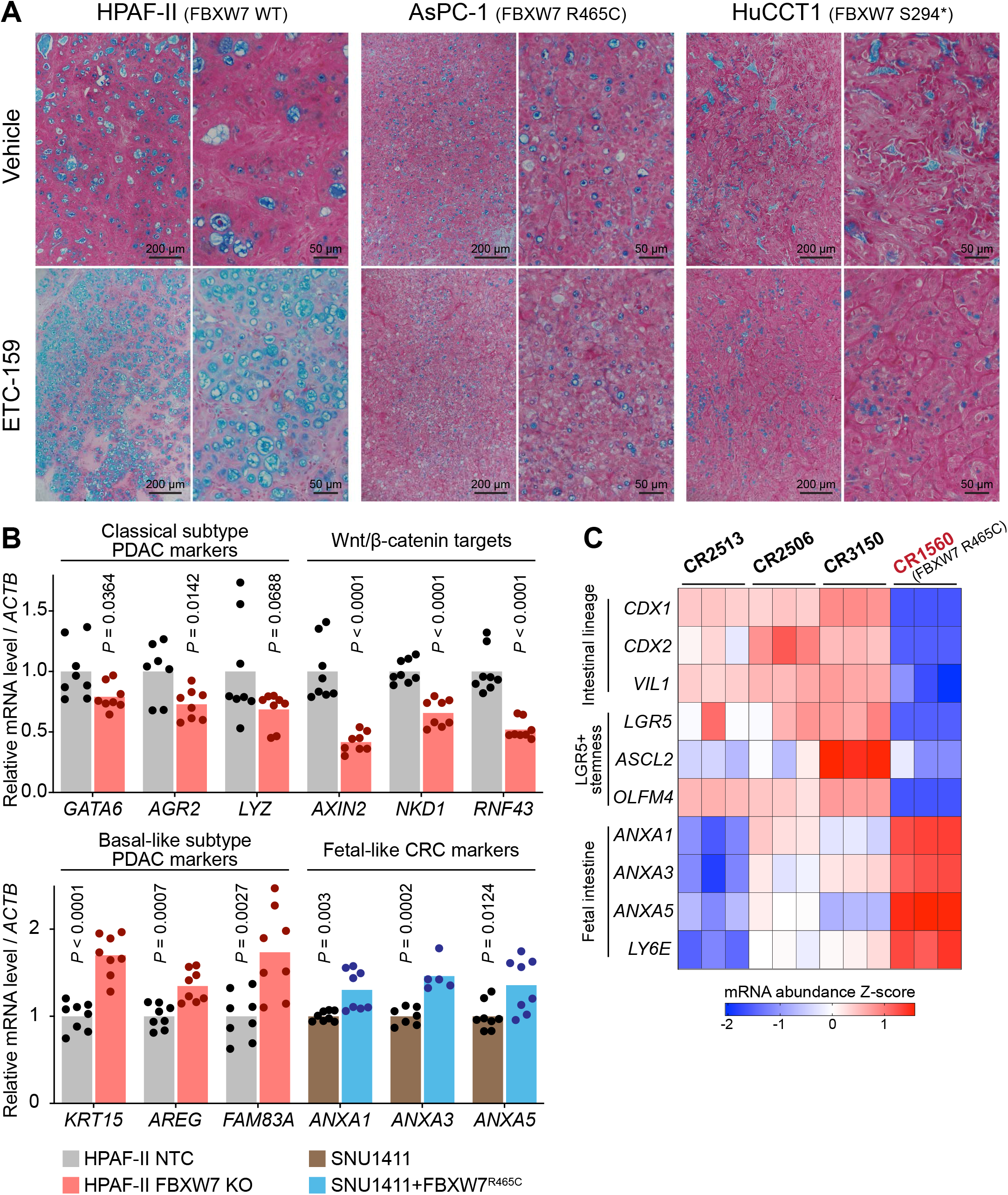
Inactivation of FBXW7 is associated with dedifferentiation and phenotypic switching in *RNF43*-mutant/*RSPO*-fusion cancers. **(A)** *RNF43*/*FBXW7*-double mutant cancer xenografts were resistant to Wnt inhibition-induced differentiation. Alcian blue (pH 2.5) mucin staining was performed on tumor samples treated with vehicle or ETC-159 **(Fig. 2B)**. **(B)** Inactivation of FBXW7 promoted molecular subtype switching in *RNF43*-mutant pancreatic tumors and *RSPO3*-fusion colorectal tumors. mRNA abundance of the indicated marker genes was measured in HPAF-II and SNU1411 xenografts. Four independent tumors per group were analyzed by RT-qPCR with technical replicates for each sample. *P* values of 2-tailed, unpaired *t* test are shown. **(C)** Among the *RSPO3*-fusion colorectal cancer PDXs **(Fig. 2C)**, the *FBXW7*-mutant PDX was dedifferentiated and showed fetal-like colorectal cancer features. mRNA abundance of the indicated marker genes was extracted from GSE73906 and shown as Z-score.

Pancreatic cancers are classified into two major subtypes, a well/moderately differentiated classical subtype and a poorly differentiated basal-like/squamous subtype^6, 33–35^. Our previous work showed that active Wnt/β-catenin signaling is present in the classical rather than the basal-like/squamous subtype^18^. Indeed, *RNF43*-mutant pancreatic tumors are enriched in the classical subtype^7, 18^. Consistent with this, the *FBXW7*-wildtype HPAF-II tumors showed clear classical subtype features^18^. However, the *FBXW7*-mutant AsPC-1 tumors showed a more mesenchymal phenotype (**Fig. 7A** and **Fig. S4**), a key feature of the basal-like subtype. This suggested a role of *FBXW7* mutation in mediating the subtype transition. To examine if changes in FBXW7 caused the subtype transition, we analyzed the expression of several subtype-related marker genes in HPAF-II tumors after FBXW7 truncation (**Fig. 7B**). Indeed, FBXW7 inactivation down-regulated the expression of the classical subtype genes and up-regulated the basal-like subtype genes. Interestingly, FBXW7 inactivation also down-regulated the expression of several well-established Wnt/β-catenin target genes (**Fig. 7B**), consistent with our previous finding that pancreatic basal-like subtype tumors have low β-catenin signaling^18^. In summary, inactivation of FBXW7 in *RNF43*-mutant pancreatic cancers leads to subtype switching.

Similarly, recent studies have divided colorectal cancer cells into two distinct cancer cell states^36–38^. One state has high LGR5+ adult intestinal stemness markers and is commonly observed in the CMS2 subtype of colorectal tumors that show high Wnt/β-catenin signaling and epithelial differentiation. The other state features a fetal-like and regenerative intestinal gene signature and is associated with tumors that have low Wnt/β-catenin signaling and mesenchymal phenotype. We found that inactivating FBXW7 by expressing a dominant-negative mutant in the *RSPO3*-fusion colorectal SNU1411 tumors up-regulated the expression of fetal-like colorectal cancer markers (**Fig. 7B**). Moreover, among the *RSPO3*-fusion colorectal cancer PDXs, the PDX harboring a *FBXW7* hotspot mutation also highly expressed the fetal-like tumor markers while had low expression of the LGR5+ stemness markers (**Fig. 7C**). These results suggest that inactivation of FBXW7 in *RSPO*-fusion colorectal cancers also leads to a subtype switching. We even noted a marked decrease of the expression of intestinal lineage markers in the *FBXW7*-mutant PDX (**Fig. 7C**), reflecting a dedifferentiated status that lacks lineage specificity.

In summary, inactivation of FBXW7 in *RNF43*-mutant/*RSPO*-fusion cancers leads to cancer cell dedifferentiation and a phenotypic switch from a Wnt-high epithelial state to a Wnt-low mesenchymal state. Previously we showed in pancreatic cancer that the classical subtype tumors require active Wnt/β-catenin signaling to antagonize the differentiation program, while the basal-like subtype tumors intrinsically inactivated the differentiation program and therefore lost β-catenin dependence^18^. Similarly, the dedifferentiated *FBXW7*-mutant tumors no longer require active Wnt/β-catenin signaling to prevent differentiation and therefore are resistant to Wnt inhibition-induced differentiation (**Fig. 7A**).

### *FBXW7*-mutant cancers are sensitive to multi-CDK inhibition

*FBXW7* mutations in *RNF43*-mutant/*RSPO*-fusion cancers lead to loss of Wnt dependence and drug resistance to anti-Wnt therapies. We examined if there are alternative strategies to treat these *FBXW7*-mutant cancers. As shown previously, Cyclin E and MYC proteins were stabilized in these *FBXW7*-mutant cancer cells and were functionally important. While it remains challenging to therapeutically target MYC directly, Cyclin E’s function can be targeted using small molecule inhibitors of the Cyclin E-coupled kinase, CDK2. As MYC may act as a general amplifier that globally enhances transcription^39, 40^, an alternative strategy of targeting MYC activity in cancer is to down-regulate the hyperactivated transcription program directly, e.g., through inhibiting the activity of transcription-associated CDKs^41^. Therefore, the goal of simultaneously targeting the Cyclin E/CDK2-driven cell cycle progression and the MYC-driven transcription in *FBXW7*-mutant cancers could be achieved by multi-CDK inhibitor treatment. To test this, we treated two *FBXW7*-mutant cancer xenograft models with the multi-CDK inhibitor dinaciclib that targets both the cell cycle-associated CDK1/2 and the transcription-associated CDK9/12^42, 43^. As a proof of concept, low-dose dinaciclib treatment significantly suppressed tumor growth in both models (**Fig. 8A**), reflecting their high sensitivity to anti-cell cycle/transcription treatment. While the dinaciclib-treated tumors were still slowly growing, perhaps due to the short *in vivo* half-life of dinaciclib (murine plasma half-life of ∼0.25 hour^42^) and low dosing frequency, these results reveal the potential of treating *FBXW7*-mutant cancers with multi-CDK inhibitors such as dinaciclib.

**Figure 8.**
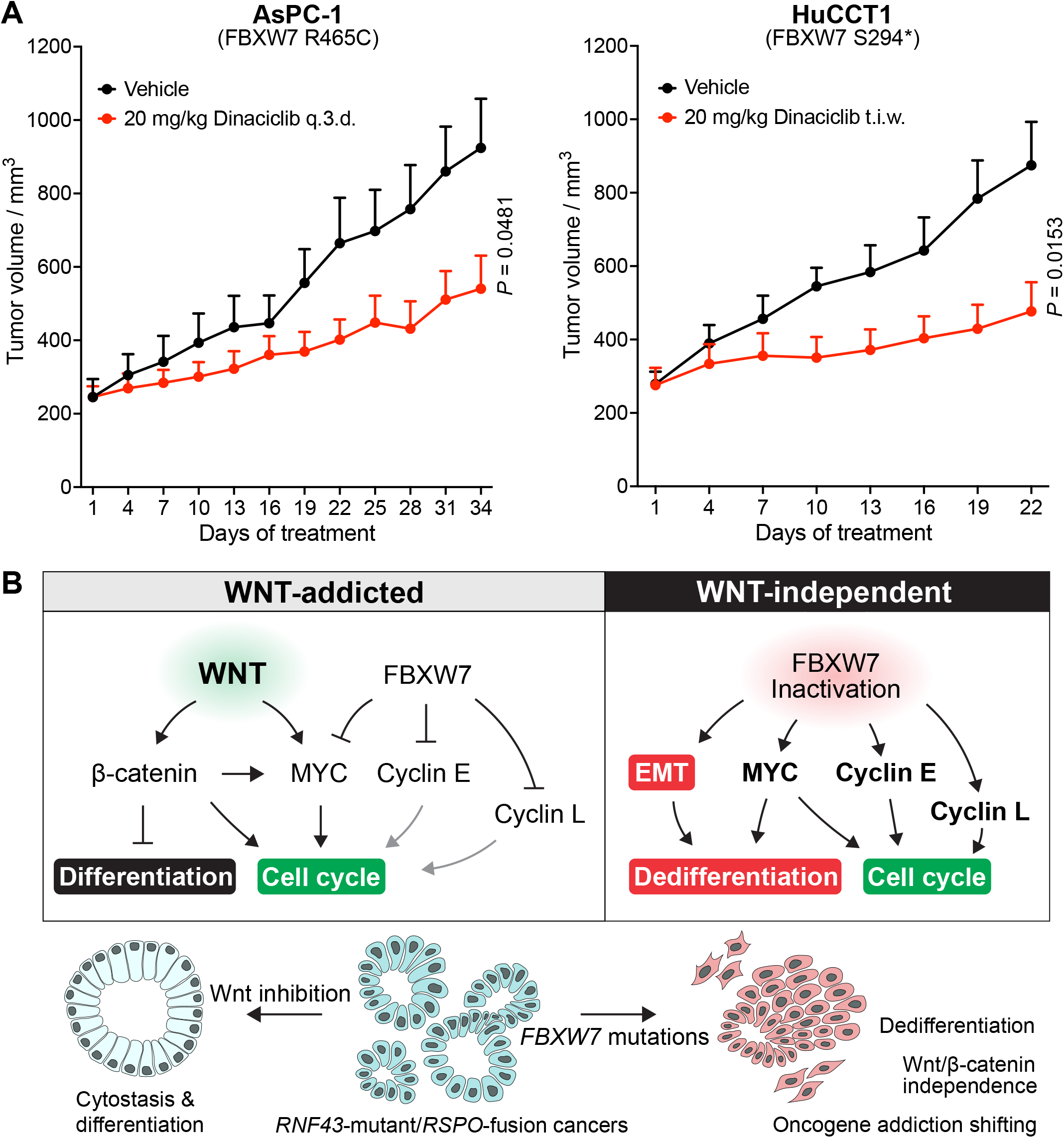
*RNF43*/*FBXW7*-double mutant tumors are sensitive to multi-CDK inhibition. **(A)** The multi-CDK inhibitor dinaciclib significantly inhibited *RNF43*/*FBXW7*-double mutant tumor growth. AsPC-1 or HuCCT1 cells were injected subcutaneously into NSG mice. After tumor establishment, mice were treated with vehicle or 20 mg/kg dinaciclib once every three days (q.3.d.) or three times a week (t.i.w.). Data are represented as mean + SEM (n = 6-8 tumors/arm). *P* values of 2-tailed, unpaired *t* test between vehicle and dinaciclib arms at the last time point are shown. **(B)** Schematic model of FBXW7 inactivation-caused Wnt independence and dedifferentiation in *RNF43*-mutant/*RSPO*-fusion cancer. Please refer to the text for detailed description.

## Discussion

Hyperactivated Wnt signaling is a potent driver of diverse human cancers, but no drug targeting the Wnt pathway has been approved for cancer therapy so far^1^. The discovery of specific *RNF43* mutations and *RSPO* fusions that caused addiction to upstream Wnt signaling in *RNF43*-mutant/*RSPO*-fusion cancers, coupled with the development of clinically tolerated upstream Wnt pathway inhibitors has generated cautious optimism for anti-Wnt therapies. However, even within this genetically defined *RNF43*-mutant/*RSPO*-fusion cancer cohort, pre-existing drug resistance represents a significant patient selection problem^16, 17^. In this study, we identified recurrent *FBXW7* mutations in *RNF43*-mutant/*RSPO*-fusion cancers and demonstrated that this is a major drug resistance mechanism.

In *RNF43*-mutant/*RSPO*-fusion cancers, a Wnt/β-catenin/MYC network maintains the active cell cycle^28^, and therefore Wnt inhibition causes cell cycle arrest followed by senescence. *FBXW7* mutations stabilize Cyclin E and MYC proteins to antagonize the cytostatic effect of Wnt inhibitors and therefore cause drug resistance (**Fig. 8B**). This is consistent with the finding that overexpression of Cyclin E or MYC in multiple other cancers, frequently due to gene amplification, is an important mechanism that confers resistance to targeted cytostatic agents, e.g., CDK4/6 inhibitors and certain tyrosine kinase inhibitors^44–49^. Notably, there might be other substrates not examined in our study that also contributed to cell cycle progression after *FBXW7* mutation. For example, another cell cycle regulator Cyclin L1 was recently identified as a novel substrate of FBXW7 and accumulated in FBXW7-inactivated cells^19^. Elevated levels of Cyclin L1 might also help to maintain the active cell cycle during Wnt inhibition.

An unexpected finding in our study that has broad implications is the loss of β-catenin dependence in these Wnt/β-catenin-driven cancers after gaining *FBXW7* mutations. We attempted to interpret this observation based on a simplified model where upstream Wnt signaling stabilizes two groups of GSK3 substrates: β-catenin and the others represented by MYC (**Fig. 8B**). In the presence of intact FBXW7 function, these Wnt-driven cancer cells require active Wnt signaling to maintain high levels of both β-catenin and MYC. After FBXW7 inactivation, activation of the MYC axis no longer requires upstream Wnt/β-catenin signaling, and additional cell cycle-promoting activities such as up-regulated Cyclin E emerge (**Fig. 8B**). We speculate that these allow the *FBXW7*-mutant cancer cells to undergo signaling and transcriptional rewiring under Wnt-low conditions that eventually makes β-catenin dispensable. Notably, the high Wnt/β-catenin activity in *RNF43*-mutant/*RSPO*-fusion cancer cells relies on sufficient paracrine and/or autocrine Wnt ligands that may come from the tumor microenvironment and may be spatiotemporally restricted. Reliance on these exogenous Wnt ligands may restrict cancer progression, e.g., from outgrowing the source of Wnts, or from metastasizing to Wnt-ligand deficient environment. Under these circumstances, *FBXW7*-mutant cancer cells that lost β-catenin dependence would gain a survival or proliferation advantage. Last but not least, inactivation of FBXW7 was recently linked to cancer immune evasion^50^, which also favors the selection of *FBXW7*-mutant cancer cells during tumor evolution.

Previously we showed in pancreatic cancer that Wnt/β-catenin signaling is important in tumors with an active intrinsic differentiation program which is reflected in the high expression of the pancreatic differentiation master regulator GATA6. The balance between Wnt signaling and the differentiation program leads to a well/moderately differentiated malignant state^18^. In pancreatic tumors whose differentiation program is intrinsically blocked, e.g., by silencing *GATA6* expression, tumors are dedifferentiated and Wnt/β-catenin signaling becomes dispensable^18^. In our current study, we observed similar dedifferentiation and phenotypic transition in *FBXW7*-mutant *RNF43*-mutant/*RSPO*-fusion cancers originated from pancreas, bile duct, and large intestine. This effect could be mediated in part by the stabilized MYC protein, as MYC overexpression has been shown to induce the expression of embryonic stem cell genes and cause the loss of tissue and lineage specificity due to dedifferentiation in cancers derived from diverse lineages^51^ (**Fig. 8B**). Moreover, several studies have shown that inactivation of FBXW7 promotes epithelial-mesenchymal transition (EMT) in multiple cancer types, although the underlying mechanisms appear to be diverse and involve the direct or indirect up-regulation of EMT factors such as Snail, Twist, or Zeb^52–55^. As EMT is an important mechanism of epithelial cell dedifferentiation, *FBXW7* mutations in *RNF43*-mutant/*RSPO*-fusion cancers may promote their dedifferentiation through inducing EMT (**Fig. 8B**), which is consistent with their mesenchymal features.

Last but not least, the *FBXW7* mutation status can serve as a biomarker to stratify *RNF43*-mutant/*RSPO*-fusion cancer patients for anti-Wnt therapies. *FBXW7* mutations may cause primary resistance to any therapy that targets the transcriptional activity of β-catenin. Notably, around one quarter of *FBXW7* mutations are missense variants of unknown significance (VUS), highlighting the importance of comprehensively characterizing these VUS. As an alternative strategy, we demonstrated that these *FBXW7*-mutant tumors were sensitive to multi-CDK inhibitor dinaciclib that targets both cell cycle and transcription. While treatment with dinaciclib alone did not achieve complete tumor growth inhibition, developing drug combinations based on dinaciclib or other multi-CDK inhibitors is a promising strategy to enhance the therapeutic efficacy. And whether such dinaciclib sensitivity is related to the *RNF43*-mutant/*RSPO*-fusion genetic background or it can be generalized to any *FBXW7*-mutant cancers deserve further investigation.

## Materials and methods

### Mutation analysis

We reviewed the literature^4, 56^ and public cancer genomics databases including cBioportal^57^, GENIE^58^, and ChimerDB4^59^ to identify *PTPRK*-*RSPO3* fusion positive colorectal tumor cases with gene mutation data available. This led to the identification of 40 non-redundant cases. Gene alterations in *FBXW7*, *APC*, *AXIN1*, or *CTNNB1* were extracted from the corresponding databases for these 40 samples.

For the *RNF43*-mutant pan-cancer analysis, 1510 *RNF43*-mutant tumor samples were identified out of 69223 non-redundant tumor cases collected in the cBioportal database. 81 samples carrying benign missense variants in *RNF43*, according to previous characterization studies, were excluded. The remaining 1429 samples were classified based on the *RNF43* mutation types. Gene alterations in *FBXW7*, *APC*, *AXIN1*, or *CTNNB1* were extracted from cBioportal for these 1429 samples.

Gene alteration information for the preclinical models used in this study was extracted from the DepMap database and selected references^26, 27, 60, 61^.

### Cell culture

The sources of cell lines used in this study are described in Supplementary Table S1. All cell lines were cultured in the corresponding culture medium recommended by the suppliers supplemented with 1% (v/v) penicillin/streptomycin (Gibco, #15140122) and in a humidified incubator with 5% CO_2_. All cell lines were regularly tested for mycoplasma contamination and confirmed to be mycoplasma free.

Lentiviruses and retroviruses were produced by co-transfecting the transfer plasmids with pMD2.G (Addgene #12259, a gift from Didier Trono) and psPAX2 (Addgene #12260, a gift from Didier Trono) or pCL-Eco (Addgene #12371, a gift from Inder Verma)^62^ into HEK293FT cells, as previously described^18^. The transfer plasmids used in this study included FuCas9Cherry (Addgene #70182, a gift from Marco Herold)^63^ for constitutively expressing Cas9 protein, Lenti-dCas9-KRAB-blast (Addgene #89567, a gift from Gary Hon)^64^ for constitutively expressing dCas9-KRAB fusion protein, FgH1tUTG (Addgene #70183, a gift from Marco Herold)^63^ for expressing doxycycline (dox) inducible sgRNA, pHR’CMV-FBXW7^R465C^ (a gift from Patrick Tan)^65^ for expressing the mutant FBXW7, MSCV MYC T58A puro (Addgene #18773, a gift from Scott Lowe)^66^ for expressing the mutant MYC, and MIGR1-Cyclin E-AA (Addgene #47498, a gift from Alex Minella)^67^ for expressing the mutant Cyclin E. After transduction, cells were selected with the corresponding antibiotics or fluorescence-activated cell sorting (FACS).

### Soft agar colony formation assay

The assay was performed as described previously^18^. Briefly, 0.6% (g/ml) agarose (Sigma, A9045) in 200 μl complete culture media was dispensed into each well of 48-well suspension culture plates to form the bottom layer. After solidification of the bottom layer, 3000 cells mixed with 0.36% (g/ml) agarose in 200 μl complete culture media were layered on top of that. After solidification of the cell/agarose layer, 450 μl complete culture media containing DMSO (0.1% v/v) or ETC-159 were added to each well. Cells were cultured for 1-2 weeks to form colonies. Colonies were stained with MTT and counted by GelCount (Oxford Optronix).

### CRISPR genome/epigenome editing

sgRNAs listed in Supplementary Table S1 were cloned into the dox-inducible sgRNA expression vector FgH1tUTG following the protocol described previously^63^. To knockout (truncate) FBXW7, HPAF-II cells stably expressing Cas9 protein were transduced with an inducible sgRNA targeting *FBXW7* (sgFBXW7#1) or a non-targeting control (sgNTC#1). Transduced cells were treated with 1 μg/ml dox for 3 days (for *in vitro* assays) or 7 days (to generate cells for injection into mice) to induce the sgRNA expression and genome editing. To analyze the genome editing, genomic DNA was extracted from the cell pellets using a salting out protocol as described previously. PCR was performed to amplify the genomic region covering the expected cutting site. The PCR amplicons were purified and subject to Sanger sequencing, and the indels were visualized using TIDE^68^.

To knockdown *CTNNB1* or *MYC*, HPAF-II, HuCCT1, or AsPC-1 cells stably expressing the dCas9-KRAB fusion protein were transduced with inducible sgRNAs targeting the promoters of *CTNNB1* or *MYC* or a non-targeting control. To examine the gene knockdown effects, transduced cells were treated with 1 μg/ml dox *in vitro* for 4 days and subject to RNA extraction and qPCR analysis. Due to the short half-life of dox at 37°C, all dox-containing media were changed every two days.

### Animal studies

Animal experiments were performed in NOD scid gamma (NSG) mice purchased from InVivos, Singapore or The Jackson Laboratory and bred and housed in specific pathogen-free conditions. For subcutaneous implantation, cancer cells were harvested from cell culture, washed with cold PBS, and resuspended in ice cold 50% Matrigel (Corning, #354248) in PBS. 5 × 10^6^ cells in 200 μl 50% Matrigel were injected into the flank of NSG mice. Tumor dimensions were measured using a caliper and tumor volumes were calculated as 0.5 × length × (width)^2^. For drug administration, ETC-159 was formulated in 50% (v/v) PEG400 in water and administered by oral gavage at a dosing volume of 5 μl/g mouse body weight following the dosing schedule indicated in each experiment. Dinaciclib was formulated in 20% (g/ml) (2-hydroxypropyl)-β-cyclodextrin in water and administered by intraperitoneal injection at a dosing volume of 5 μl/g mouse body weight following the dosing schedule indicated in each experiment. To induce the sgRNA expression *in vivo*, mice were fed *ad libitum* with standard diet containing 600 mg doxycycline/kg (Specialty Feeds, SF08-026) until the end of the study. The tumors and organs with metastases, if any, were harvested and snap-frozen in liquid nitrogen followed by storage at -80 °C, or immediately fixed in 10% neutral buffered formalin for two days at room temperature followed by paraffin embedding. The SingHealth Institutional Animal Care and Use Committee approved all animal studies, which compiled with applicable regulations.

### Western blotting

Fresh cells or snap-frozen tumor tissues were homogenized in 4% SDS in water using a tissue homogenizer. Protein concentration of the lysate was measured by the bicinchoninic acid (BCA) assay. Equal amounts of total protein of each sample were resolved by 10% SDS-PAGE followed by wet transfer to PVDF membranes. Western blotting was performed following standard methods. Primary and secondary antibodies used in this study are described in Supplementary Table S1. The SuperSignal West Dura Extended Duration Substrate (Thermo Scientific, #34075) or Cytiva Amersham ECL Select Western Blotting Detection Reagent (Fischer Scientific, #45000999) were used for developing the chemiluminescence blots. The images were captured digitally using the Bio-Rad ChemiDoc Touch Imaging system or the LI-COR Odyssey Imaging system.

### Histological analysis

Formalin-fixed paraffin-embedded tissue sections were deparaffinized and rehydrated. Hematoxylin and Eosin (H&E) staining was performed with standard methods. Immunohistochemistry (IHC) for β-catenin, GFP, and Ki67 was performed as described previously^18^. Antibodies used for IHC are described in Supplementary Table S1. Alcian blue (pH 2.5) mucin staining was performed using the Alcian Blue Stain Kit (Abcam, ab150662). Bright field images were taken with a Nikon ECLIPSE Ni-E microscope.

### RNA extraction and quantitative PCR (qPCR)

Total RNA from fresh cells or snap-frozen tumor tissues was extracted using the QIAGEN RNeasy Kit. cDNA was synthesized using the MiRXES BlitzAmp cDNA synthesis kit. Real time qPCR was performed using the Bio-Rad SsoAdvanced Universal SYBR Green Supermix on Bio-Rad CFX96 or CFX384 Touch real-time PCR machines. Primer sequences are described in Supplementary Table S1. All qPCR primers are human specific.

### Gene set enrichment analysis (GSEA)

Transcriptomic data of the *RSPO3*-fusion colorectal PDXs treated with RSPO3-neutralizing antibody or control were extracted from GSE73906^15^. Gene set enrichment analysis was performed using the GSEA software (www.gsea-msigdb.org)69.

### Statistical analysis

Statistical analyses were performed using GraphPad Prism 10 or as otherwise stated. Two-tailed unpaired *t* tests were used to analyze data involving direct comparison between an experimental group and a control group. *P* value < 0.05 or FDR < 0.1 were considered to be statistically significant.

## Author contributions

ZZ and DMV designed the study. ZZ performed the experiments and analyses. DMV supervised the study. ZZ and DMV wrote the manuscript.

## Acknowledgements

We gratefully acknowledge Yunka Wong and Xinang Cao for their invaluable technical supports and advice. We thank members of the Virshup lab for useful discussions. ETC-159 was a generous gift from the Experimental Drug Development Centre of Singapore. This research was supported in part by the National Research Foundation Singapore and administered by the Ministry of Health’s National Medical Research Council under Singapore Translational Research (STaR) Award MOH-000155 (to DMV). Figure 5A was created with BioRender.com.

## Notes

**Conflict of interest:** David M. Virshup has a financial interest in ETC-159.

### Competing Interest Statement

David M. Virshup has a financial interest in ETC-159

### Summary of Updates

Updated acknowledgments.

