## Supplemental Figures for "Recurrent *FBXW7* mutations bypass Wnt/β-catenin addiction in cancer"

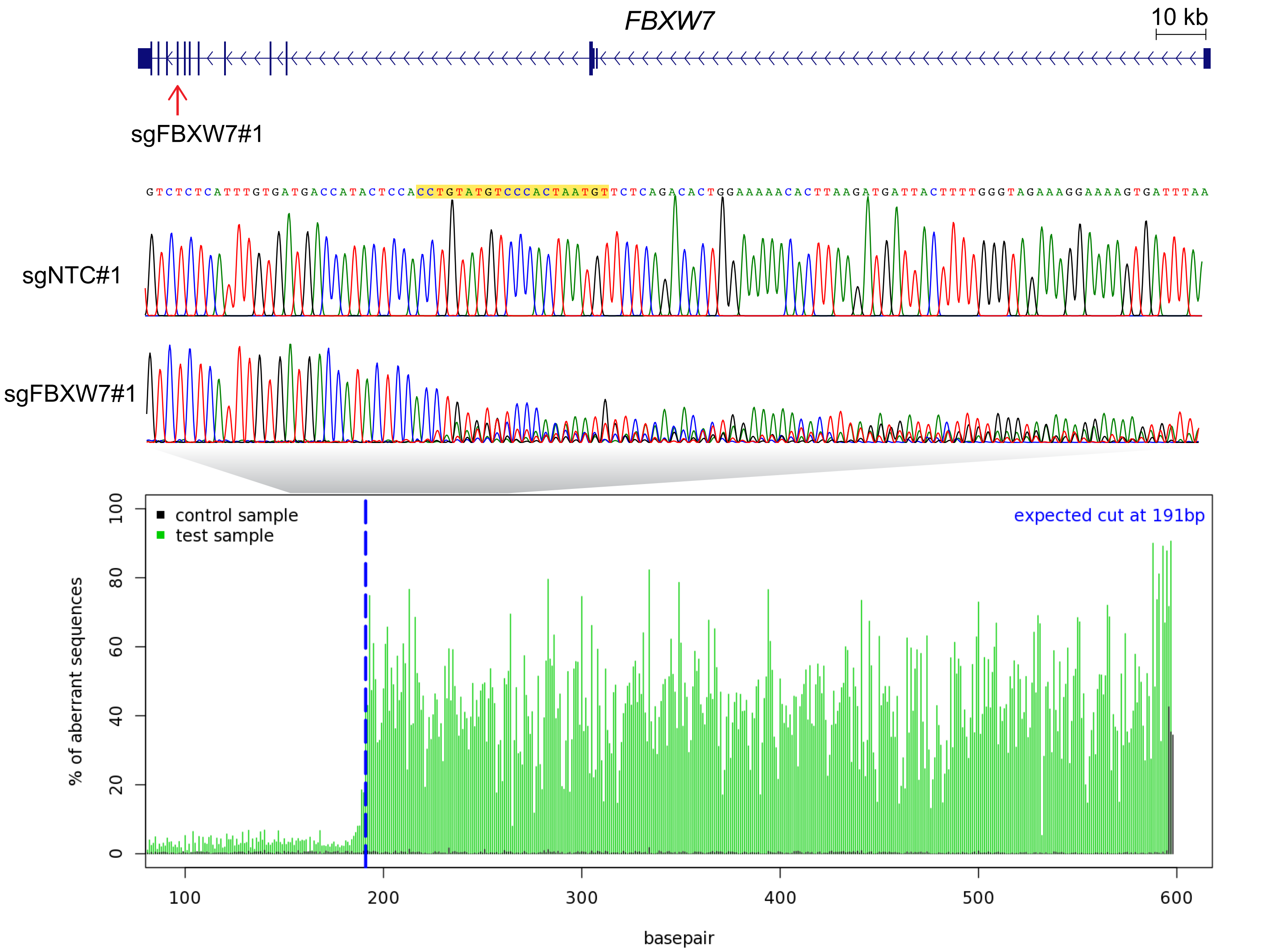


**Supplementary Figure S1. CRISPR/Cas9-mediated truncation of FBXW7 in HPAF-II cells.** The genome position targeted by the sgRNA (sgFBXW7#1) is shown on the top. This sgRNA led to considerable indels at the expected cutting site on the genome (corresponding to FBXW7 amino acid position 420) compared to the non-targeting control sgRNA (sgNTC#1). The reference DNA sequence is shown and the sgRNA sequence is highlighted in yellow. Comparison of the Sanger sequencing signals visualized using TIDE is shown at the bottom. Please refer to the Materials and methods part for experimental details.

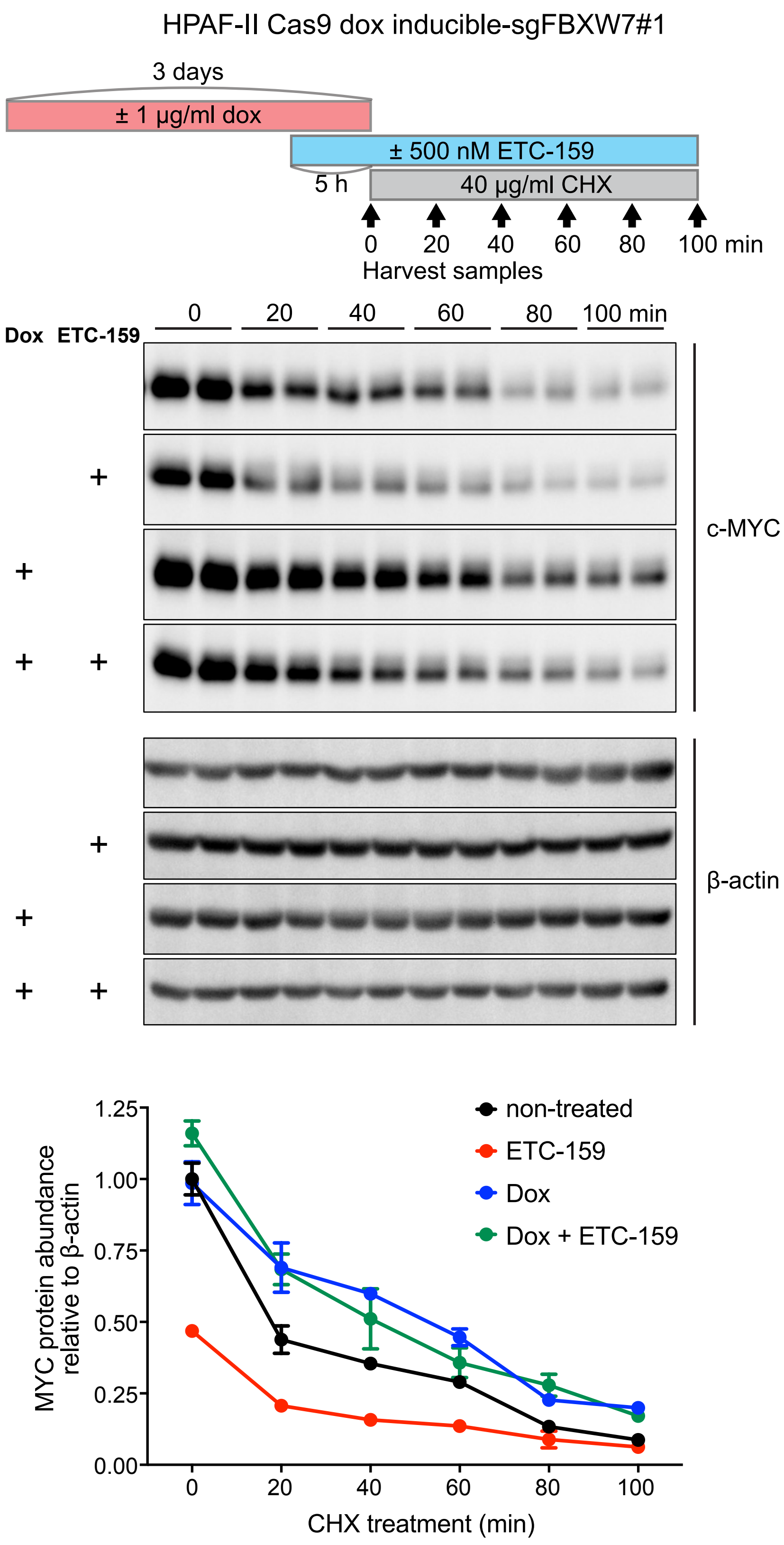


**Supplementary Figure S2. Truncating FBXW7 in HPAF-II cells stabilized MYC protein in a cycloheximide (CHX) chase assay.** HPAF-II cells constitutively expressing Cas9 and carrying a doxycycline (dox) inducible FBXW7-targeting sgRNA (sgFBXW7#1) were treated with dox, the PORCN inhibitor ETC-159, and CHX as indicated. Cell lysates were analyzed by western blotting for MYC. β-actin was used as a loading control. n = 2 biological replicates/condition. Quantification of the band intensity normalized to the time 0 non-treated condition is shown at the bottom.

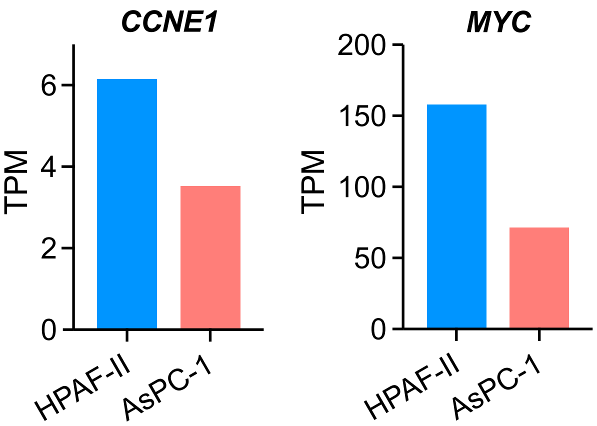


**Supplementary Figure S3. mRNA abundance of *CCNE1* and *MYC* in HPAF-II and AsPC-1 cells.** Gene expression data were extracted from the DepMap database.

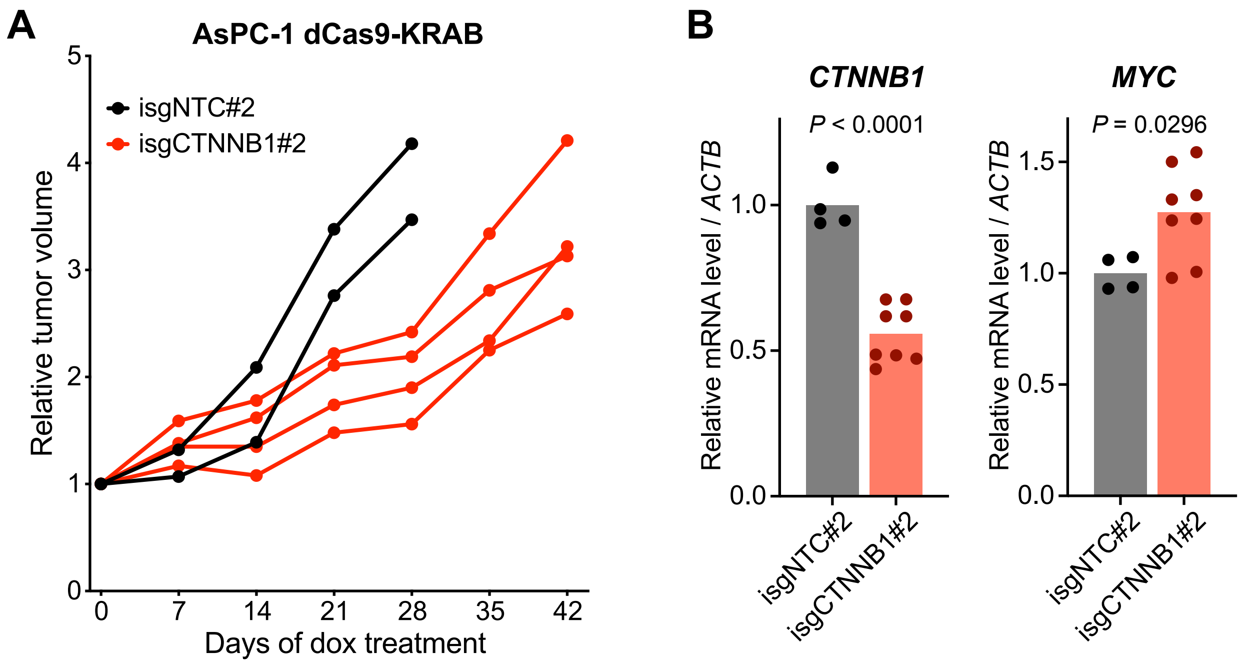


**Supplementary Figure S4. Knockdown of *CTNNB1* in *FBXW7*-mutant AsPC-1 xenografts.** **(A)** AsPC-1 cells stably expressing dCas9-KRAB were transduced with a doxycycline inducible sgRNA targeting *CTNNB1* promoter (isgCTNNB1#2) or a non-targeting control (isgNTC#2). These cells were subcutaneously injected into NSG mice. After tumor establishment, mice were put on continuous doxycycline feed till the end of the study. The individual tumor growth curves are shown. Note the increase in tumor growth rate after day 28 in the *CTNNB1* targeted tumors. **(B)** Relative mRNA abundance of *CTNNB1* and *MYC* in AsPC-1 tumors harvested in the end of the study. Tumors from two isgNTC#2 tumors and four isgCTNNB1#2 tumors were analyzed by RT-qPCR, with technical replicates for each sample. *P* values of unpaired t test are shown.

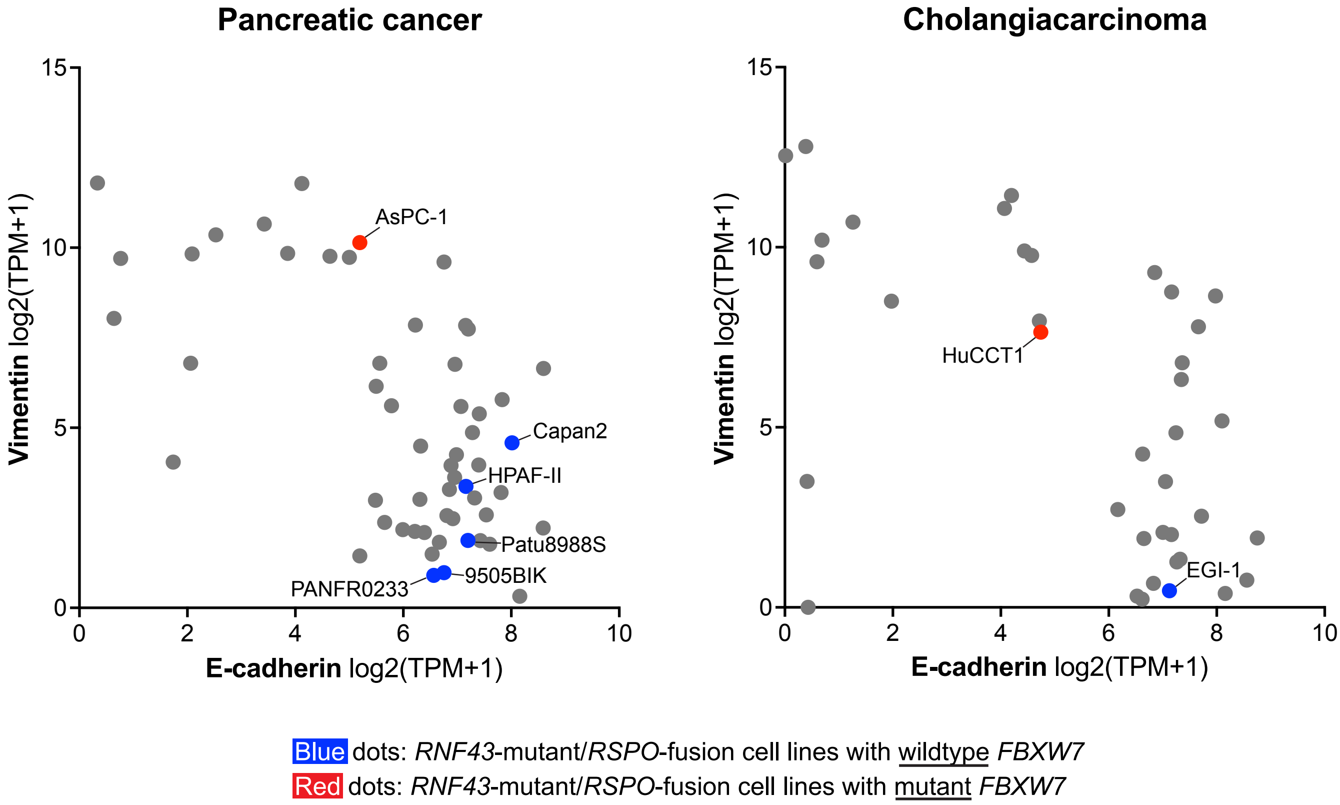


**Supplementary Figure S5. *FBXW7*-mutant *RNF43*-mutant/*RSPO*-fusion pancreatobiliary cancer cell lines show mesenchymal features.** mRNA abundance of the epithelial marker E-cadherin and the mesenchymal marker vimentin in pancreatic cancer cell lines (left) and cholangiocarcinoma cell lines (right) are shown. Gene expression data were extracted from the DepMap database. Each dot represents an individual cell line.
